# Core-2 *O*-glycans are required for galectin-3 interaction with the osteoarthritis related protein lubricin

**DOI:** 10.1101/2019.12.20.884148

**Authors:** Sarah A. Flowers, Kristina A. Thomsson, Liaqat Ali, Shan Huang, Yolanda Mthembu, Suresh C. Regmi, Jan Holgersson, Tannin A. Schmidt, Ola Rolfson, Lena I Björkman, Martina Sundqvist, Anna Karlsson, Gregory D. Jay, Thomas Eisler, Roman Krawetz, Niclas G. Karlsson

## Abstract

Synovial fluid lubricin (proteoglycan 4) is a mucin-type *O*-linked glycosylated (60% of the mass) biological lubricant involved in osteoarthritis (OA) development. Lubricin has been reported to be cross-linked by synovial galectin-3 on the lubricating articular surface. Here, we confirm that binding to galectin-3 depended on core-2 *O*-linked glycans, where surface plasmon resonance of a recombinant lubricin (rhPRG4) devoid of core-2 structures lacked binding capacity to recombinant galectin-3. Both galectin-3 levels and interactions with synovial lubricin were found to be decreased in late-stage OA patients coinciding with an increase of truncated and less sialylated core 1 O-glycans. These data suggest a defect cross-linking of surface active molecules in OA and provides novel insights into OA molecular pathology.

## INTRODUCTION

Lubricin is a large glycoprotein encoded by the *Prg4* gene, which is found in synovial fluid (SF). Lubricin is produced by articular chondrocytes and synoviocytes in the cartilage surface layer, and secreted into the SF. In healthy joints, lubricin molecules are bound to the surface of articular cartilage, are chondroprotective, and provides lubrication at very low friction under high stress^1, 2^. The lubricating properties of lubricin are dependent, in part, by dense *O*-glycosylation^3^.

Lubricin has been associated with pathological conditions in the joint tissue, highlighted by the fact that the mutations of the *Prg4* gene is associated with the camptodactyly-arthropathy-coxa vara-pericarditis syndrome (CACP)^4^, a rare arthritis like autosomal recessive disorder causing joint abnormalities. The lubrication properties of lubricin also suggest that the protein is associated with arthritic diseases, since it has been proposed that defect lubrication should aggravate joint degradation^5^. Osteoarthritis (OA) is the dominating arthritic disease with high prevalence in elderly people and involves cartilage degradation in the articular joints, leading to pain and restricted motion^6^. Boundary lubrication of OA SF deficient in lubricin has been shown to be lowered in *in vitro* experiments^7^.

Different isoforms of lubricin are found throughout the body. In addition to SF and the boundary articular cartilage, it has also been detected in low amounts in menisci, blood, urine, and tendons^8, 9 10 11, 12^. A possible immunological role of lubricin has been suggested, since it has been shown to be involved in sepsis in a mouse model^13^, being the most upregulated protein detected in hepatic tissue. Lubricin is also associated with the plasma membrane of human neutrophils^14^, and a lubricating and protective function of Lubricin has been reported in ocular surfaces^15 16^.

The full-length isoform found in SF consists of 1,404 amino acids (aa) equivalent to 150 kDa, but gains in molecular mass to approximately 300 kDa when fully *O*-glycosylated. The protein contains a central mucin domain, consisting of 59 imperfect repeats of the aa sequence ‘EPAPTTPK’, where the threonine residues are potential sites for diverse and dense *O*-glycosylation. Despite its gene name, it is not a typical proteoglycan, and has merely a single proposed glycosaminoglycan site that may or may not be present/occupied^17^. The mucin domain is flanked by two somatomedin B-like domains on the *N*-terminal side, and a hemopexin domain on the *C*-terminal side, domains which have been proposed to interact with extracellular matrix (ECM) proteins such as cartilage oligomeric matrix protein (COMP), collagen II and fibronectin^18 19^. The glycans on the lubricin mucin domain are the main contributors to its molecular weight and are responsible for the lubricating function of the molecule, attracting water to the superficial layer and providing repulsive negative charges to the cartilage superficial layer.

Recombinant human lubricin (rhPRG4) has been evaluated for a large number of potential clinical applications as a friction reducing lubricant on biological surfaces^16^. Forms of rhPRG4 have been investigated for use as an agent to prevent abdominal adhesion, treatment of bladder impermeability, as a lubricant supplement in contact lens hydrogels and as a therapeutic in patients suffering from dry eye disease^20 21, 22, 23^. In rodent models, addition of lubricin has been shown to delay the progression of OA^24^. Intra-articular injections of rhPRG4 in guinea pigs^25^ and Yucatan minipigs^25^ with medial meniscal destabilization, a model system for posttraumatic OA, exhibited decreased cartilage damage and inflammation. Lubricin may therefore be a potential candidate for treatment of OA.

Galectins are glycan-binding proteins, with many proposed intra- and extra-cellular functions within cell regulation, the immune system and cancer progression^26^. They bind a large range of β-galactosides present both on *N*- and *O*-linked glycoproteins, with binding affinity highly dependent on branching and terminal residues^27^. Galectin-3 has been shown to be associated with chondrocytes and proposed to be involved in osteoarthritis pathogenesis^28, 29^. It is unique among the galectins, as it is the only galectin that comprise a C-terminal carbohydrate recognition domain linked to a non-lectin N-terminal collagen-like domain making it possible for galectin-3 protein subunits to oligomerize^30, 31^. In the joint, galectin-3 together with lubricin have been proposed to generate a complex stable lubricating network on the articular cartilage^32^. Limited studies have addressed galectin-3 binding to *O*-glycans. Galectin-3 displays only a weak binding affinity to the T-antigen (Galβ1-3GalNAc) on synthetic MUC1 glycopeptides^33, 34^, which is an highly abundant *O*-glycan on lubricin, and proposed to be responsible for galectin-3 lubricin interaction^32^.

Since the carbohydrates play such a crucial role in the function of lubricin, we decided to use state-of-the-art glycomics and glycoproteomics to analyse the differences in glycoforms of synovial lubricin from OA patients and control individuals, and to compare these with a type of rhPRG4 expressed in CHO-cells. We confirmed the presence of galectin-3 in synovial fluids, and explored differences in the binding of galectin-3 to the various *O*-glycoforms. Our results enable us to correlate pathological glycosylation changes in OA with a potential functional defect superficial layer, and provides leads into recombinant lubricin glycodesign for optimal galectin-3 binding ability.

## RESULTS

### DECREASED CORE-2 AND SIALYLATION ON SF LUBRICIN FROM OA PATIENTS

We wanted to address if an alteration of the *O*-linked glycosylation of synovial lubricin could be associated with OA pathology. Detailed quantitative analysis of the released *O*-glycans was performed. Lubricin isolated from SF of controls (n = 7) was compared to lubricin isolated from late stage OA patients (n = 7). *O*-linked oligosaccharides from lubricin in age- and sex-matched OA patients’ and controls’ SF were released by reductive β-elimination and analyzed with quantitative MRM on a QTRAP 6500 triple quad-linear ion trap hybrid mass spectrometer as has been described in detail previously^35^. The MRM method was developed to semi-quantify a range of twenty-two core-1 and −2 and −3 glycans on native lubricin from both controls and OA patients^35^. Although the abundances of these glycans were found to differ between the two sample types, the glycan repertoire was the same (Fig. 1). A significant increase of the unmodified core-1 structure (T-antigen) on lubricin isolated from OA patients (40% of all glycans) was detected compared to normal controls (30% of all glycans). This increase in glycan truncation was accompanied by a significant decrease in a range of lower abundant structures, predominantly, core-2 sialylated and sulfated structures. Altogether, this resulted in a reduction in charged structures in the OA samples (57% of all glycans) compared to the controls (67% of all glycans). A 45% reduction of core 2 structures in OA lubricin (total of 6.4% core-2 glycans) compared to control lubricin (3.5% core-2 glycans) was also detected. This indicated that changes in *O*-glycosylation, and specifically the loss of sialylation and core-2 as seen here, is associated with OA.

**Figure 1.**
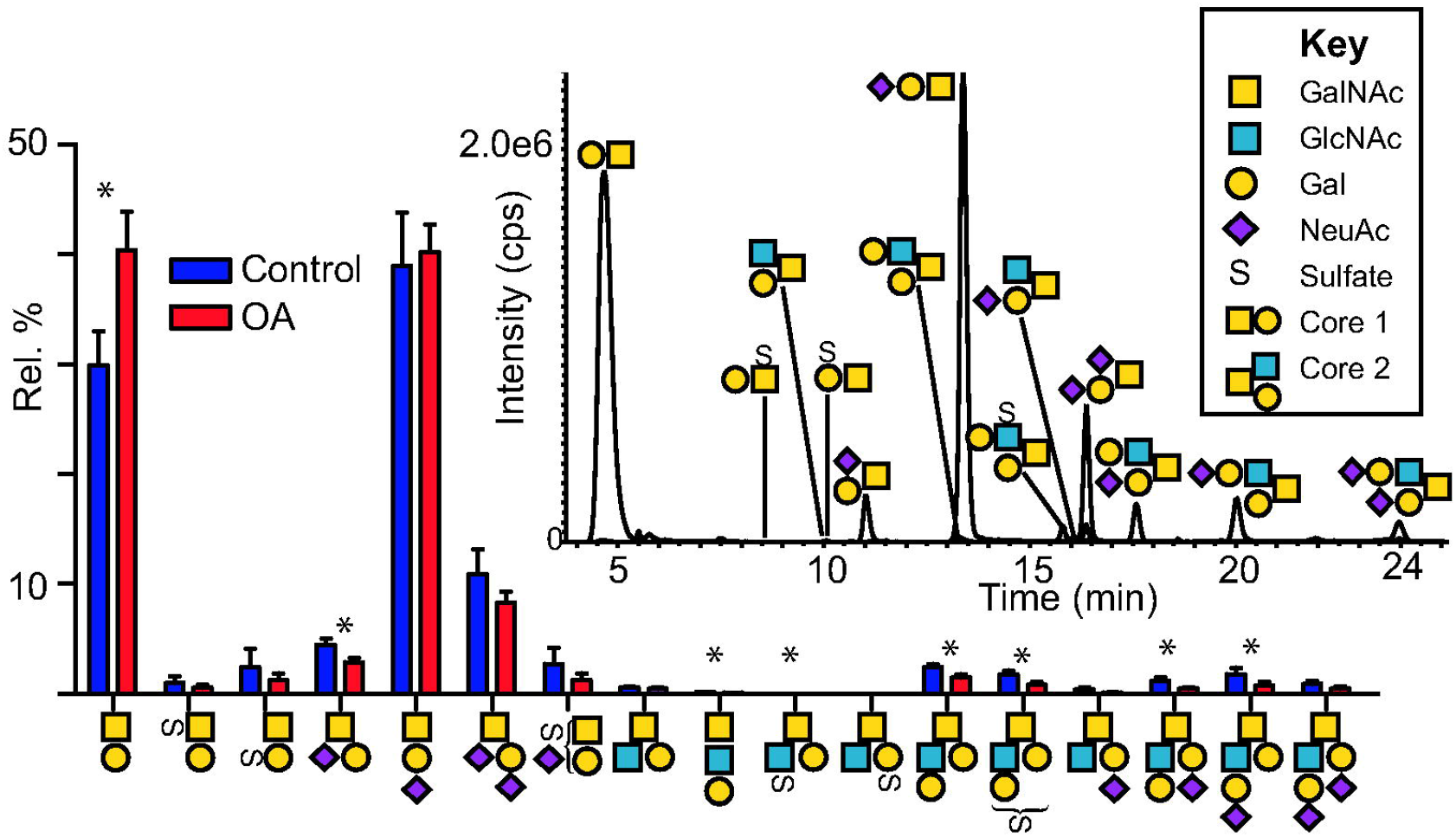
MRM analyses of *O*-glycans from SF lubricin purified from control and OA individuals. Relative quantification comparison of control individuals (n = 7) and OA patients (n = 7) released glycans from SF lubricin showing core 1 and core 2 *O*-glycans with the most dominant structures being the unmodified and monosialylated core 1. Graph shows average and standard deviation. Inset, example of extracted ion chromatogram of *O*-glycans from lubricin from a control individual. *p ≤ 0.05. Symbol key is according to the SNFG nomenclature for glycans. Statistical analyses, two-tailed Student’s *t*-tests, were carried out to compare the normal and OA synovial lubricin glycan relative abundances.

### CHO EXPRESSED rhPRG4 HAS MAINLY SIALYLATED CORE 1 O-GLYCANS

Given the potential of rhPRG4 as a biopharmaceutical lubricant where the function relies on the *O*-linked glycosylation, we undertook the task to analyze the glycosylation of CHO expressed rhPRG4 to identify the presence of core-1, core-2 glycans and sialylation and compare it to *O*-glycosylation of the human SF lubricin.

The composition of *O*-glycans released from rhPRG4 was analyzed using the same complete MRM lubricin method as for human lubricin. These analyses were performed on two different batches of rhPRG4, which revealed a composition solely of core 1 structures (Fig. 2). The most abundant oligosaccharide was the sialylated core 1 NeuAcα2-3Galβ1-3GalNAc structure, which made up 85% of all glycans. A low proportion of neutral *O*-glycans (12% of all glycans) and a higher amount of charged glycans (88% of all glycans) were found in this rhPRG4. The glycosylation pattern of rhPRG4 is similar to what has been reported for CHO cell *O*-glycosylation previously, with the abundance of core 1 structures specifically the linear monosialylated NeuAcα2-3Galβ1-3GalNAc structure^36^.

**Figure 2.**
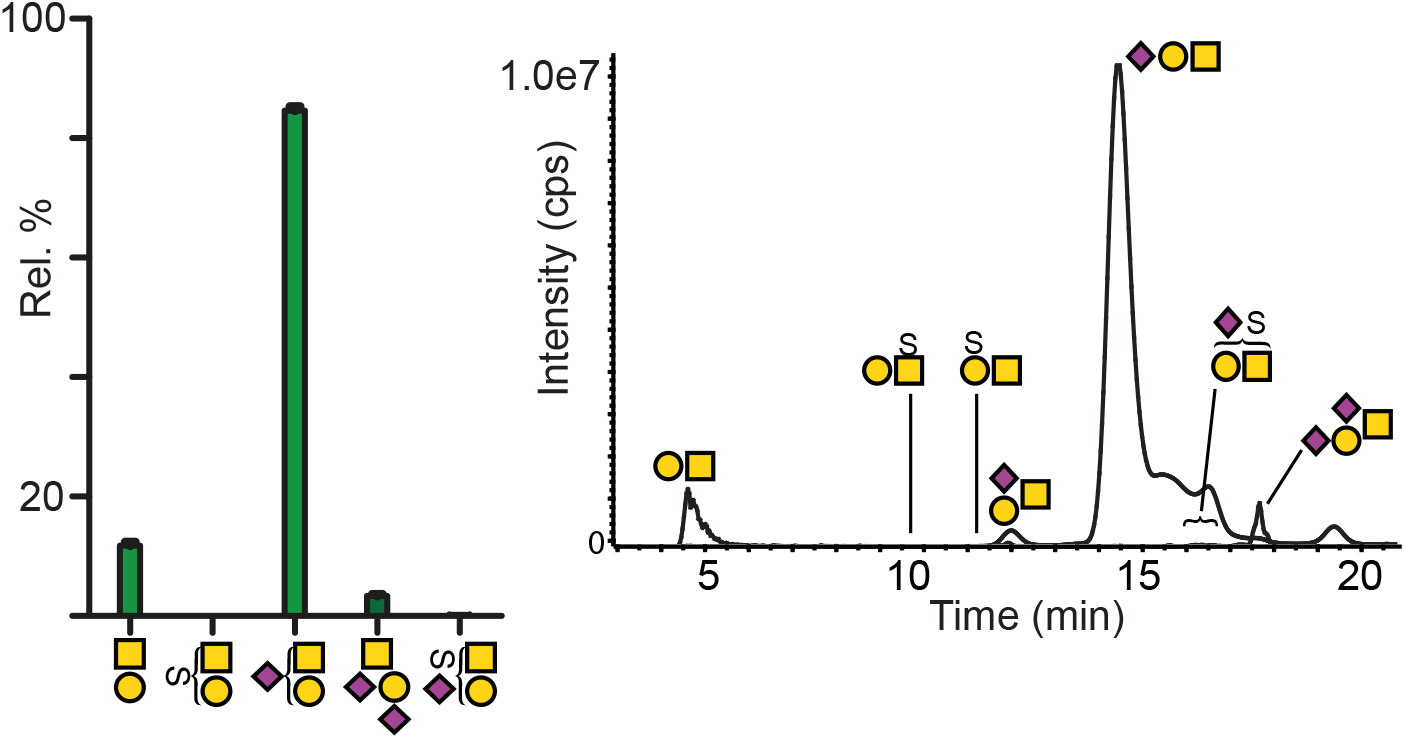
Relative quantification by MRM of released *O*-glycans from rhPRG4 showing only core 1 structures and an abundance of the monosialylated structure. Data is from two batches each repeated in duplicate. Graph shows average and standard deviation. Inset, example of extracted ion chromatogram of rPRG4. For key of symbols, see Fig. 1.

### GLYCOPROTEOMICS REVEAL LOCATION OF TN, SIALYL-TN AND CORE TYPE 1 STRUCTURES BOTH WITHIN AND OUTSIDE rhPRG4 MUCIN DOMAIN

We next performed a glycoproteomics study of rhPRG4 (with and without partial deglycosylation) to identify glycosylated aa sites, and compared with glycosites previously reported for native human lubricin from pooled SF from OA and Rheumatoid Arthritis (RA) patients^37^. In all, we detected 318 glycopeptides from rhPRG4 covering 172 glycosylated Ser/Thr sites, the monosaccharide compositions consistent with core 1 type O-glycans GalNAcα1-(Tn-antigen), Galβ1-3GalNAcα1-(T-antigen), NeuAcα2-3Galβ1-3GalNAcα1-(sialyl T) and NeuAcα2-3Galβ1-3(NeuAcα2-6)GalNAcα1-(disialylated T). The results are summarized in Fig. 3 and in Supplementary Table 1. Glycopeptide coverage in rhPRG4 was found to be similar to native human lubricin, with the majority of the glycopeptides originating from the S/T rich mucin domain (aa 232-1056)(Fig. 3a). Site specificity overlapped well with native lubricin and within the mucin domain (Fig. 3b, c and Supplementary Fig. 1). 83 sites were detected in rhPRG4 which have not been identified in native lubricin (Fig. 3b). The majority of the glycopeptides contained 1-3 glycans, although up to five glycosylation sites were detected on one peptide (<u>TT</u>PE<u>TTT</u>AAPK, aa sequence 926-936). Glycopeptides generated after partial deglycosylation revealed additional site-specific information., enabling the detection of larger glycopeptides in the size range of 10-20 aas. In all, 31 additional glycosylated Ser/Thr were identified after partial deglycosylation (Supplementary Table 1). Approximately 77% (132 of the 172) of the glycosylated Ser/Thr were found in peptides originating from the mucin like ‘repeat’ region (aa 348-855, Fig. 4a) of rhPRG4. The most abundant aa sequence in this region was the ‘EPAPTTPK’, repeated 17 times in the molecule. Thirteen ‘EPAPTTPK’ derived glycopeptides were detected, glycosylated on either or both Thr residues (Supplementary Table 1). We found 19 glycosylation sites in the *N*-terminal region (aa 125-347) and 26 in the *C*-terminal region (aa 1057-1404). Most of these sites (16 and 25 sites from the *N*- and *C*-terminal region, respectively) were found in the non-repeat regions in the STP rich domain flanking the mucin domain 348-1057 (Supplementary Fig. 1). The most intense glycopeptides identified in rhPRG4 correspond well with those identified from pooled SF from patients with RA and OA. The main difference was that in regions with clustered Thr and to some extent also Ser, we usually found that more sites were positively identified in rhPRG4 compared to native lubricin.

**Figure 3.**
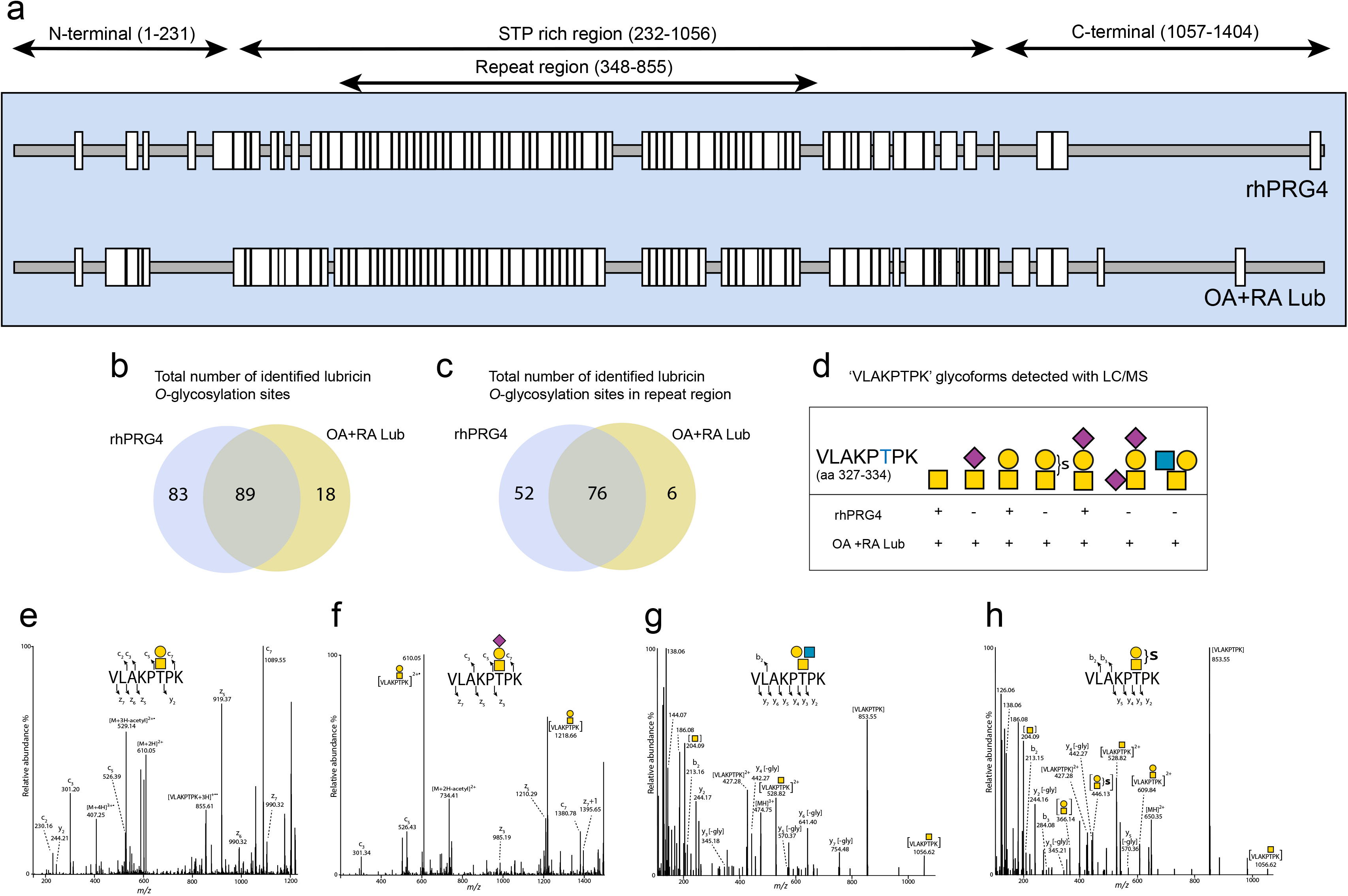
O-linked glycosite map of rhPRG4 compared to synovial lubricin. (A), glycopeptides from recombinant (rhPRG4) (this report) and SF lubricin pooled from five patients with OA and RA (this report and ref^37^) were analyzed with LC/MS with CID and ETD fragmentation before and after partial deglycosylation with sialidase and *O*-glycanase. (B) Total number of *O*-glycosylation sites confirmed by ETD in rhPRG4 and from previous analysis of SF lubricin from pooled RA and OA patients on whole lubricin and (C) in the repeat region (aa 348-855) (D) Inserted table shows the glycovariant of the glycopeptide VLAKPTPK found in SF lubricin and in rhPRG4. (E-G) Examples MS/MS of the glycopeptides variants of ‘VLAKPTPK’ from lubricin identified from rhPRG4 (ETD) and from an OA patient (HCD); *m/z* at 406.90 (3+) from rhPRG4 interpreted as a T-antigen (Galβ1-3GalNAcol) (e); *m/z* at 503.93(3+) from rhPRG4 interpreted as sialyl-T (NeuAcα2-3Galβ1-3GalNAcol) (f) *m/z* 474.59 (3+) interpreted as a core 2 glycan (Galβ1-3[GlcNAcβ1-6]GalNAcol) from an OA patient (g), *m/z* 649.83 (2+) interpreted as a sulfated core 1 type glycan (HSO_3_ + Galβ1-3GalNAcol) from an OA patient (h). For key of symbols, see Fig. 1.

**Figure 4.**
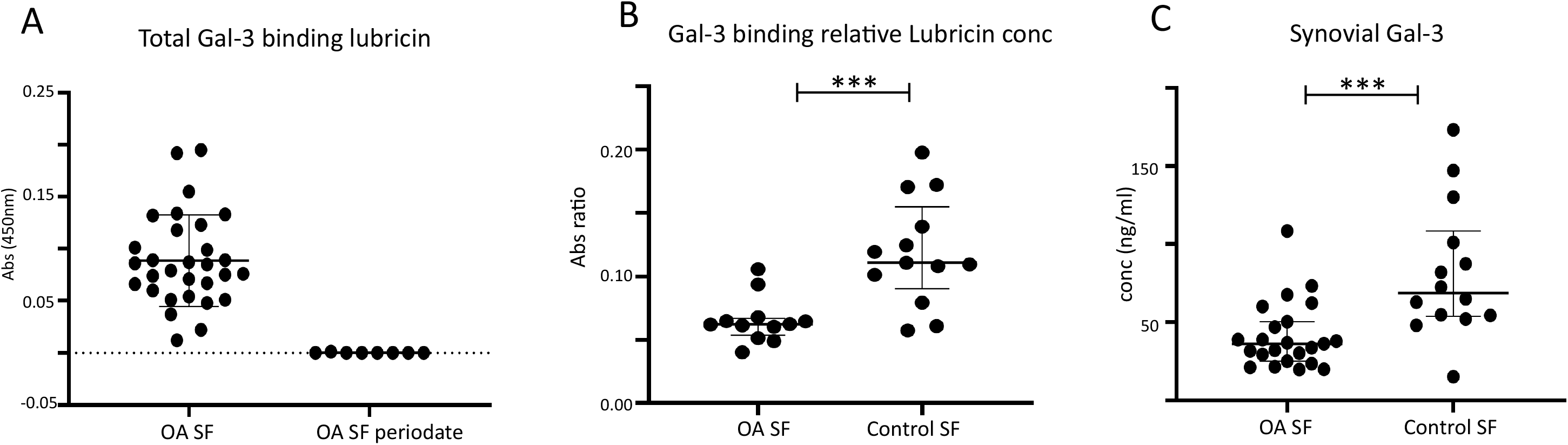
Lubricin *O*-glycans and Galectin-3. (A) Recombinant galectin-3 binding to OA lubricin reflecting both variation in lubricin concentration and glycosylation. Galectin-3 binding to lubricin from SF of late stage OA patients (n=30) was measured using a sandwich lectin ELISA assay (see Materials and Methods). Treatment of OA SF (n=8) with periodic acid shows that the binding to galectin-3 was glycosylation dependent. Data is represented as median with interquartile range, and significant difference was calculated by non-parametric Mann-Whitney test with ***p<0.001. (B) Galectin-3 binding of late stage OA patients and control SF lubricin normalized to the synovial lubricin level. Galectin-3 and relative lubricin concentration was measured by ELISA on 12 OA and 13 control SF samples, and ratio of absorbances at 450 nm were calculated for each sample. Samples were measure in duplicates. Data was represented as median with interquartile range, and significant difference was calculated by non-parametric Mann-Whitney test with ***p<0.001. (C) Endogenous galectin-3 in SF. Galectin-3 concentrations in SF were measured in samples from 23 late stage OA patients and 14 controls. Data was represented as median with interquartile range, and significant difference was calculated by non-parametric Mann-Whitney test with ***p<0.001.

### GLYCOPROTEOMICS OF SF LUBRICIN FROM OA PATIENTS

Since previous glycopeptide analyses was performed on pooled SF samples^37^, we pursued more sensitive MS analyses using HCD-MS and MS/MS, addressing the characterisation of SF lubricin glycopeptides from two individual OA patients. This approach revealed 171 and 158 lubricin derived glycopeptides from the two patients, respectively, and the full glycopeptide list is found in Supplementary Table 1. Glycopeptide coverage did largely match that found in the samples derived from rhPRG4 and our previous study of pooled native lubricin from OA and RA patients (Fig. 3a and Supplementary Table 1), suggesting that many glycosylation sites coincide.

To illustrate the type of information obtained from SF lubricin versus rhPRG4, seven glycopeptides with the peptide backbone sequence ‘VLAKPTPK’ (aa 327-334, Fig. 3d) were found. This peptide backbone sequence contains only one potential *O*-glycosylation site (T). In addition to simple core 1 structures identified in rhPRG4 (Fig. 3e, f), we also detected more exotic type *O*-linked glycans attached to native lubricin. Examples of two of these spectra are shown in Fig. 3g, h. Both spectra were dominated by a major fragment ion corresponding to the peptide backbone without its glycosylation (*m/z* 853.55). The core 2 glycopeptide variant of this glycopeptide was identified containing two HexNAc and one Hex, consistent with a (Galβ1-3(GlcNAcβ1-6)GalNAcα1-core 2 sequence. Low molecular oxonium ions at *m/z* 138, 144, 168, 186, and 204 originated from fragmentation of monosaccharide residues and were diagnostic for glycan containing peptides. The other example from SF lubricin is a ‘VLAKPTPK’ glycopeptide with a T-antigen with an associated sulfate residue (Fig. 3h). We have previously shown that sulfated core 1 *O*-glycans exist on SF lubricin both as 3-linked and 6-linked sulfate to the non-reducing end Gal residue, but also to the 6-position of the reducing end GalNAc residue^38^. The diagnostic glyco oxonium ions at *m/z* 204, 366 and 446 were glycan fragment ions consisting of HexNAc, HexNAc-Hex and HexNAc-Hex + sulfate, respectively. This peptide example illustrates that sulfoglycopeptides can successfully be identified amongst other glycopeptides in positive ion mode without targeted approaches that has been suggested^39^.

One *N*-glycosylated site was found on lubricin from both recombinant and OA patient derived lubricin at Asn1159. The two dominant glycoforms found in both samples were of high mannose type, Hex_5_HexNAc_2_ and Hex_6_HexNAc_2_ respectively. Annotated HCD and CID spectra of these are displayed in Supplementary Fig. 2, 3.

### DECREASED GALECTIN-3 BINDING TO SF LUBRICIN AND DECREASED ENDOGENOUS GALECTIN-3 IN SF OA

We investigated galectin-3 and its binding lubricin in SF from OA patients. The experiment was performed in SF from 30 OA patients using recombinant galectin-3 in an ELISA assay. We found that the level of galectin-3 active lubricin present in SF varied approximately 1-2 orders of magnitude between the individuals (Fig. 4a). Periodate treatment, which destroys the glycans abolished the binding, proving that the interaction was purely dependent on glycosylation. We then normalized the amount of galectin-3 binding lubricin with the amount of lubricin in each sample. This approach allowed the galectin-3 binding capacity of lubricin in SF to be assessed and compared to that of control individuals. We could show that lubricin from late stage OA (n=12) was significantly less able to bind galectin-3 compared to the controls (n=13) (p<0.001) with a factor of almost 2 (Fig. 4b). Furthermore, we generated data that showed a significant decrease (50%) in the total amount of endogenous galectin-3 in SF of the late stage OA (n=23) compared to controls (n=14) (Fig. 4c). These data suggest the level of galectin-3 and its ability to bind SF lubricin, are altered as a consequence of OA.

### GALECTIN-3 AND CORE 2 GLYCOSYLATION

A potential functional impact of the altered OA glycosylation identified here was investigated further. Lubricin glycans are essential for several critical properties of lubricin involving lubrication and binding properties and has been suggested to bind to galectin-3 to improve lubrication of the joint via the abundant T-antigen epitopes (Galβ1-3GalNAcα-)^32^. However, we observed decreased lubricin binding to galectin-3 in OA in parallel with an increase of this glycan on OA lubricin, which lead us to test the hypothesis that the lubricin galectin-3 interaction may be mediated by epitopes presented on extended core 2 glycans.

To address if core 2 glycans compared to the simpler core 1 glycans were responsible for the galectin-3 binding we used recombinant glycotechnology. We modified the CHO-cell core-1 glycosylation by transfection with the human core-2 GlcNAcβ1-6 glycosyltransferase^40^. A reporter *O*-linked mucin type glycoprotein (PSGL-1/mIgG2b) expressed in these cells was shown to be able to bind recombinant galectin-3 (Fig. 5a, b). Without transfection of the glycosyltransferase, no binding to PSGL-1/mIgG2b was observed. This demonstrated that core-2 type oligosaccharides are key for recognition by galectin-3.

**Figure 5.**
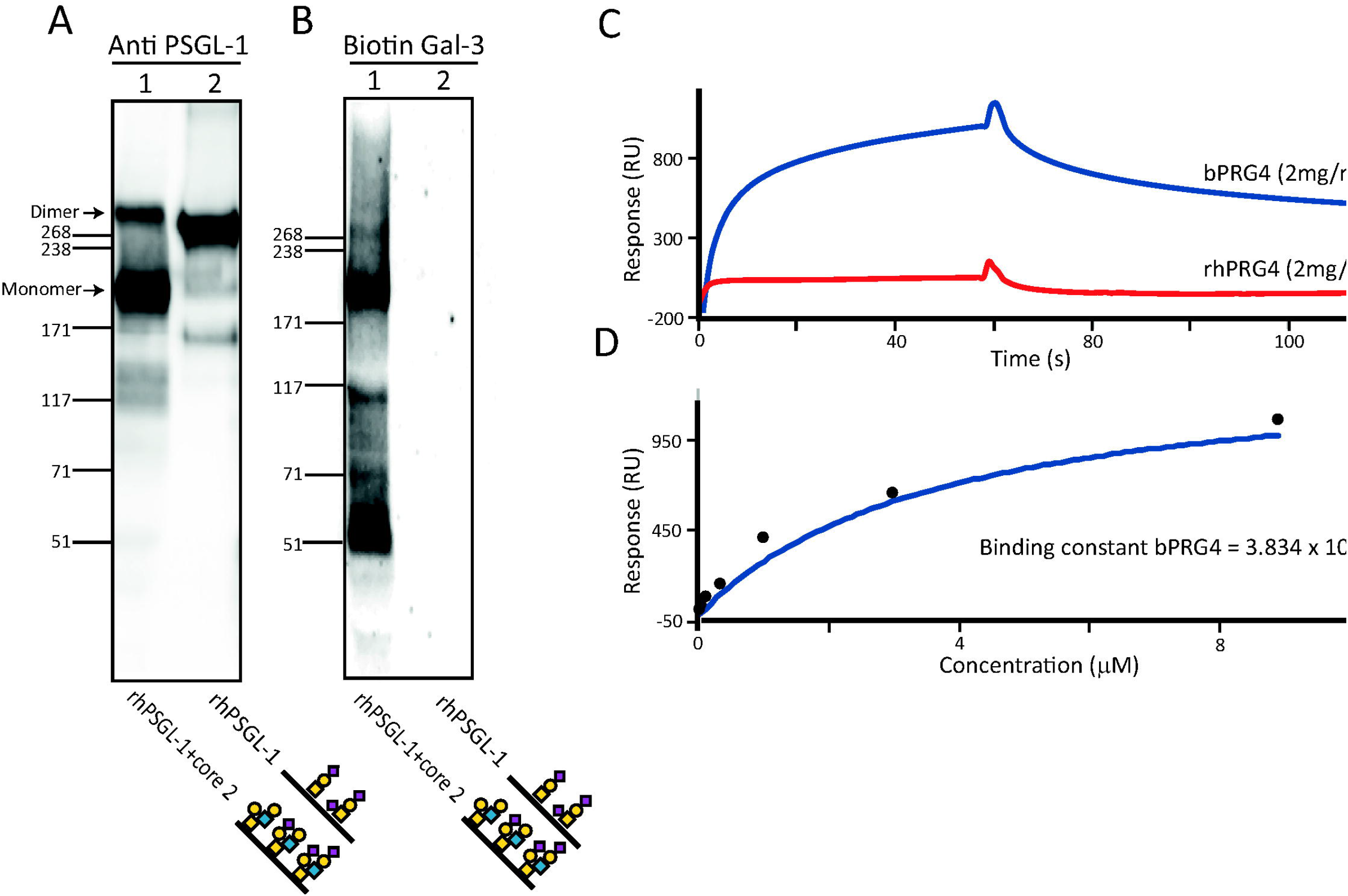
Western blot analysis of immune purified recombinant PSGL-1/mIgG2b expressed in CHO-K1 cells with and without co-expression of core 2 glycosyltransferase (β1,6-N-acetylglucosaminyltransferase 1). PSGL-1 (3 μg/lane, (A); 12 μg/lane, (B)) was analyzed under non-reducing conditions by SDS-PAGE, and developed with western blot probed with anti-PSGL1 (A) and recombinant biotinylated galectin-3 (biotin-Gal 3) (B). The pictures show dominant *O*-glycans found on PSGL-1/mIgG2b, (with core 1 glycans) and PSGL-1/mIgG2b+core 2 (with core 2 glycans) and have been described elsewhere^40^. (C) Surface plasmon resonance (SPR) experiment showing binding of bovine lubricin (bPRG4) to recombinant galectin-3, while rhPRG4 did not bind. (D) Calculation of binding constant of bPRG4/Galectin-3 from SPR based on molecular mass of bPRG4 estimated to 220 kDa. The pictures show prominent O-glycans on the two proteins, with symbol key displayed in Fig. 1.

**Figure 6.**
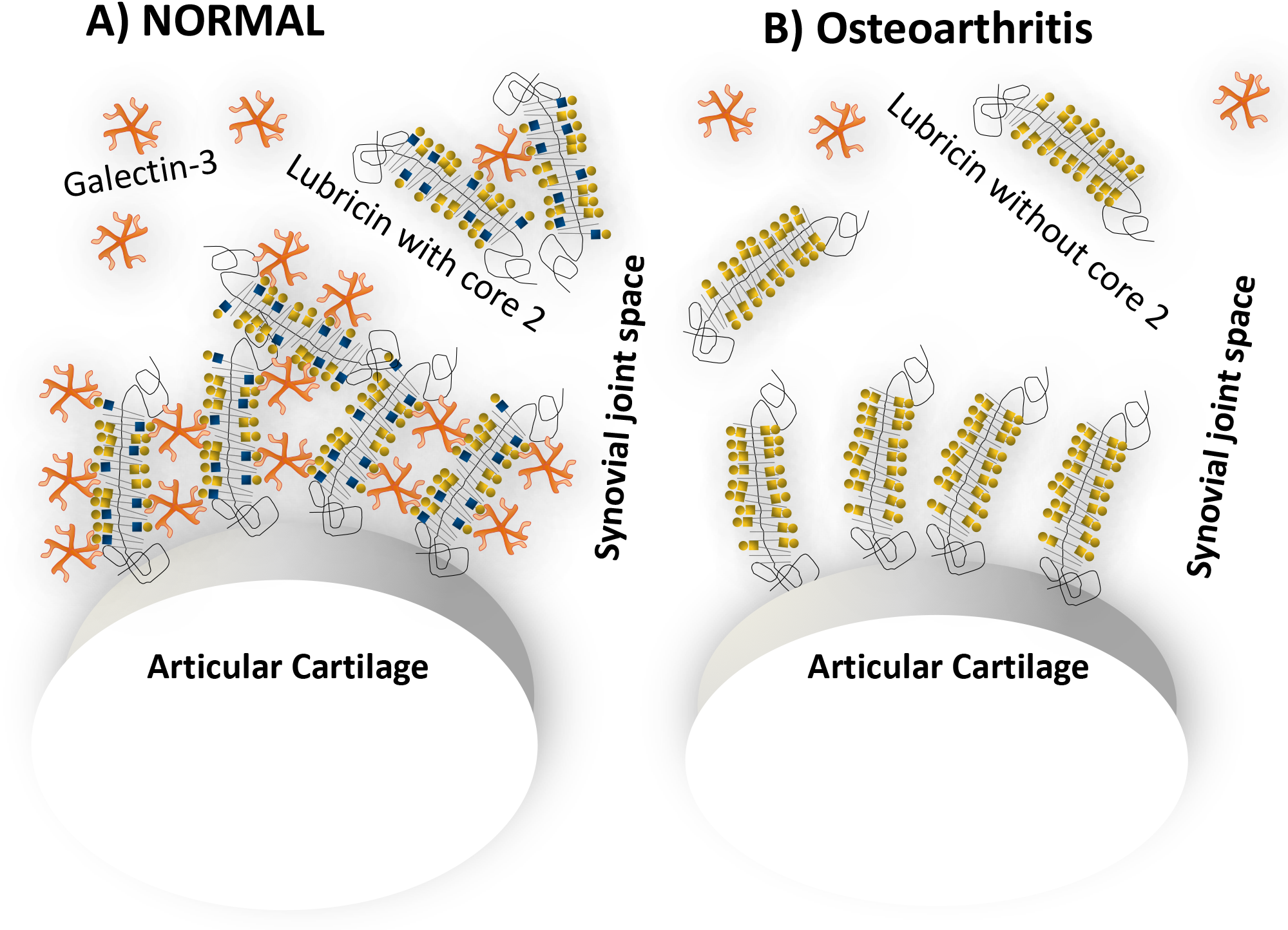
Proposed model for the organization of lubricin and galectin-3 in the superficial articular layer in normal joint and in OA. (A) In the normal joint, Synovial lubricin adheres to the cartilage surface and contains significant amount of LacNAc epitopes (represented as ■○) that allows galection-3 interaction and crosslinking of lubricin molecules. (B) In OA, lubricin is still capable of adhering to the cartilage and provide boundary lubrication, but the decreased amount of galectin-3 and its decreased interaction with lubricin destabilize the glycosylated articular surface.

Using surface plasmon resonance (SPR), the core-1 dominant CHO-rhPRG4 glycosylation was not able to bind galectin-3, while the positive control of lubricin isolated from bovine chondrocyte cell culture with both core 1 and core 2 glycosylation (Supplementary Table 2) displayed significant interaction with galectin-3 with a K_D_ calculated to 3.8 μM (Fig 5c,d). The data together indicates that the avidity of the multimeric galectin-3 relies on numerous epitopes within lubricin and its mucin domain. The PSGL-1/mIgG2b *O*-linked oligosaccharides generated after transfection of CHO-cells using human core-2 GlcNAcβ1-6 glycosyltransferase terminate with sialic acid after adding a LacNAc unit to the natively expressed core 1 structures^40^. This is also the type of core-2 structures found on lubricin^35^.

## DISCUSSION

We show that lubricin from SF of control and OA patients display differences in *O*-glycosylation. OA patients have higher levels of T-antigen (Galβ1-3GalNAcα-) compared to the sialylated form (NeuAcα2-3Galβ1-3GalNAcα-), and also a relative decrease of low abundant core 2 glycans. We also undertook an extensive analysis of a type of CHO-derived recombinant lubricin, with respect to the relative amounts of *O*-glycans as well as site occupancy on the aa level in the protein backbone. We investigated the hypothesis that the core 2 *O*-glycans on lubricin may provide better binding sites than core 1 *O*-glycans for synovial multimeric galectin-3, a binding which has been proposed to stabilize the synovial lubricating boundary layer in the healthy joints. In accordance with this, we showed that SF lubricin from OA patients displayed less binding to galectin-3 compared to controls.

### Truncation of SF lubricin O-glycosylation in OA and galectin-3

The MRM analyses of the released *O*-glycans from SF lubricin from controls and OA patients demonstrate a 10% increase of the truncated unsialylated core 1 structures (Galβ1-3GalNAcα1-) in late stage OA, accompanied by a decrease a low abundant core 2 glycans (Fig. 1). It has been demonstrated previously that the high number of negatively charged sialic acid residues located as they are within the center of the lubricin molecule with positively charged lysine and arginine rich *N*-and *C*-termini, creates the amphoteric nature of the lubricin glycoprotein^37^. The glycosylation, along with the organization of sialic acid within the molecule, has been associated with an effective lubrication property of lubricin^37, 41^. This organization creates a central hydration shell and sparingly glycosylated *N*- and *C*-termini, available to bind to other cartilage proteins including COMP and collagen II. An effective boundary lubrication model with firm adherence to the articular cartilage surface of glycomediated lubrication^18, 19, 37^ can be proposed. The reduction in sialylation in OA observed in this report can be predicted to have an effect on the overall charge and potentially the lubricating properties of the lubricin molecule in OA.

Our data on SF lubricin O-glycan truncation in OA also bears implications on binding to galectin-3. Previous studies have indicated that galectin-3 exhibits only weak affinity to *O*-glycoproteins and *O*-glycopeptides expressing single core 1 O-glycans (Galβ1-3GalNAc or NeuAcα2-3Galβ1-3GalNAc)^34, 42^. Instead, internal LacNAc glycans on extended oligosaccharides have been described to be the high affinity ligands for galectin-3^43^. Here, we suggest that interaction between galectin-3 and native lubricin is mediated by short single LacNAc extended core 2 structures. Indeed, expression of the core2β-1,6-N-acetylglucosaminyltransferase (C2GnT) responsible for biosynthesis of core-2 O-glycans has been shown to enable binding of galectin-3 to bladder tumor cells^44^.

rhPRG4 has shown to have similar lubrication properties as bPRG4 *in-vitro*^45^. Hence, the proposed Galectin-3 binding to lubricin^32^ are likely to only fine-tune the boundary lubricating glycosurface of the joint. In regards of galectin-3 level in SF, it has also been shown that patients with RA have increased level of galectin-3^46, 47, 48^, while the literature is less conclusive about the situation in OA. Our data shows that compared to the normal SF, the galectin-3 levels are diminished in late stage OA. Our hypothesis included that an altered ability of OA lubricin to bind galectin-3 together with a decreased galectin-3 levels in OA may contribute to destabilization of the boundary lubrication and contribute to joint degradation (Fig. 5). However, this notion needs to be reconciled with results from a large animal OA model where CHO cell derived rhPRG4 was shown to retard cartilage degeneration^25^. Our data shows that glycobiology hold clues for unravelling the multifaceted enigmatic OA aetiology and pathology. Of note, this is even without taking into account that cartilage harbors the largest volume of glycans (keratan- and chondroitin sulfate) in the body.

### Glycomap of native lubricin

A previous map of native lubricin was performed on tryptic and Lys-C digested *O*-glycopeptides of intact and partially de-glycosylated glycopeptides from a pooled sample^37^. Here, we included sites identified on native lubricin from individual patients using a fast scanning Orbitrap mass spectrometer to make the glycosite map more complete. Due to fast scanning and sensitivity, low abundant glycoforms could be selected for MS/MS, without any prefractionation and multiple digestion strategies. We identified 171 and 158 glycopeptides from the two lubricin tryptic digests from the OA patients, many which has not been detected before. This sensitive approach allowed the first detection of a sulfated *O*-glycopeptide from lubricin (Fig. 4, 5), and evidence for an *N*-glycosylation site on lubricin (Supplementary Fig. 1, 2). In our former study of pooled lubricin from both OA and RA patients, we reported 185 *O*-glycopeptides, however after excluding those glycopeptides that were enzymatically de-glycosylated, this number dropped down to 122. However, partial deglycosylation may still need to be carried out to identify sites within the mucin domain. For example, the glycopeptides with aa sequences ‘EPAPTTTK’, ‘KPAPTTPK’ and ‘SAPTTPK’, which are found repeated 8, 4, and 4 times in the protein respectively, were either not detected or poorly represented in the analyses of individual OA patients, but were found in both rhPRG4 and in our previous study after deglycosylation. In this paper we identified eight sites in native lubricin in addition to already recorded sites^37^.

### The rhPRG4 molecule

The analyses here showed the detailed glycomic and glycoproteomic analyses of the heavily *O*-glycosylated CHO expressed protein lubricin giving glycosite-specific information as well as detailed *O*-glycan identification and quantification. Obtained glycopeptides from rhPRG4 digested with trypsin or Lys-C, were analyzed with LC/MS/MS and CID/ETD fragmentation before and after partial deglycosylation. In total,172 glycosylation sites (Ser/Thr) were identified compared to the previous reported 107 for native lubricin obtained with the same approach^37^. The majority of the glycosylation sites of rhPRG4 were found on peptides that were repeated between 2-17 times in lubricin mucin domain. The dominant glycans found corresponded to simple core 1 type *O*-glycosylation found on this protein whereas native lubricin also contained core 2. The difference in glycosylation sites is probably a reflection of differences in specificity and level of GalNAcα1-polypeptide transferases. Human synovial cells are dominated by T1-, T2-, T5- and T15-GalNAc transferase^37^, while CHO cell glycosylation is dominated with the hamster equivalent of human T2-transferase^49^. We may also attribute differences in detected glycosylation sites to the increased glycosylation complexity of native lubricin compared to this rhPRG4 and; the decreased complexity in glycans in this rhPRG4 would cause decreased glycopeptide diversity leading to increased molar amounts and detectability of individual glycopeptides. In addition, the differences in the level of glycosylation could also contribute to differences in efficiency in the digestion of the mucin domain.

Our glycoproteomic data shows that glycans on rhPRG4 are positioned in the same central domain as in synovial lubricin. Lubricin (both native and recombinant) contains one of the largest mucin domains where glycosylation sites have been mapped. In the synovial lubricin, 37% of Ser/Thr-residues were glycosylated in the STP rich region aa 232-1056, whereas in rhPRG4, 60% of Ser/Thr-residues were found to be glycosylated. However, we need to state that the site localization relies on positive identification, so absence of detection does not mean that remaining sites are not glycosylated.

Difference in glycosylation between native lubricin and rhPRG4 invoke a question about the impact of glycosylation on the *N*-and *C*-termini involved in interactions. Lubricin has for instance been shown to bind non-covalently to COMP within aa105-202 and covalently by specific disulfide bonds between aa64-86^19^.Glycosylation sites were identified just adjacent or within these regions (Supplementary Table 2)

The *O*-glycomic data shows that rhPRG4 glycosylation is dominated by the linear monosialylated core 1 structure, similar to native synovial lubricin. rhPRG4 as such contains the main molecular functions attributed to native lubricin. Although there is a reduction in disialylated structures, there is an overall increase in sialylation giving the rhPRG4 molecule a more negatively charged central domain similar to SF lubricin from control individuals rather than lubricin found on OA patients (Fig. 1, 2). However, rhPRG4 does have a more limited range of glycans with no core 2 or extended core 1 structures present. These structures may be involved in other glycan-glycan interactions with the ECM, interacting with galectin-3 or involved in the immune system^14^. Overall, it is clear that both the type and the specific pattern of glycosylation are quality attributes to the lubrication and binding functioning of this very heavily glycosylated protein. The data indicates that glycodesign could help to further improve functionality of rhPRG4. Active modulation of *O*-linked glycosylation for biologics is an unexplored area. As of yet, it is unknown whether these changes of lubricin in SF are deleterious to the function of the molecule *in vivo*, or whether alterations in lubricin glycosylation are a biological response to OA conditions, where the change in negative charges is an attempt to alter the lubricating function of the expressed lubricin, a metabolic response to simplify and maintain expression of a less energetically expensive molecule in the face of exogenous stress, or to potentiate a downstream signaling through altered galectin-3 release. Given the complexity of timing and severity impacts on biological functions within the OA joint, in can be summarized that there are important alterations in the amount and type of glycosylations in a critical lubricating system, and that functional and inflammatory aspects of the disease are likely regulated by the expressed sugars, all of which warrant additional research and focus for heretofore unexplored pathogenic mechanisms in a very serious disease.

## METHODS

### Samples, human tissues and cells

Recombinant human Lubricin (rhPRG4) expressed from CHO cells were purified as described elsewhere^2, 50^. Bovine lubricin (bPRG4) was purified using CsCl gradient and anion exchange chromatography, and purity confirmed with SDS-PAGE/western blot and mass spectrometry as described elsewhere^51^. SF samples from OA patients that were scheduled for total knee replacement (TKR) (n=2, age 76 and 84) for the glycopeptide analyses were collected during therapeutic joint aspiration at Sahlgrenska University Hospital (Gothenburg, Sweden). SF samples for *O*-glycomic analyses were collected from patients with symptomatic chronic knee OA requiring aspiration (OA patients’ (n = 7, average age 56 y, range 34-72 y). The patients for *O*-glycomics were diagnosed as having knee OA by 2 sports medicine physicians following a review of the patient’s symptoms, a physical examination, and plain-film radiography. Control samples (n = 7, average age 56 y, range 31-72 y) were collected post mortem within 4 hr. Samples were collected from the knee by aspiration and clarified (3,000 g for 30 min at 4°C) and stored at −80°C. SF samples for galectin-3 studies were obtained from late-stage OA patients prior to TKR, mean age 69 y (n=30, range 58-78 y) at Sahlgrenska University Hospital and at Danderyds Hospital (Stockholm, Sweden). Control SF samples was collected as described above, mean age 65 years (n=18, range 47-76 years old). All samples were collected after written consent and the procedure was approved by the ethics committees at Sahlgrenska University Hospital, and University of Calgary Human Research Ethics Board. Recombinant galectin-3 used in all experiments unless stated elsewise were provided by Professor Haakon Leffler, Lund University, Sweden.

### Release and LC-MRM analysis of oligosaccharides from rhPRG4 and synovial lubricin

Lubricin from SF of OA patients (n=7) and controls (n=7) were isolated via a multistage process. The acidic SF proteins were isolated from SF samples by DEAE chromatography as previously described^52^. Samples were reduced and alkylated and separated by SDS-PAGE using NuPAGE^®^ 3-8% Tris-acetate gels to allow isolation of the lubricin. Gels were transferred to PVDF membrane by semi-dry transfer blotting, stained with Alcian blue and destained with methanol^53^. The lubricin band was excised from each sample individually and the *O*-linked oligosaccharides were released by reductive β-elimination (50 mM sodium hydroxide, 0.5 M sodium borohydride) overnight followed by cleanup on cation exchange columns (AG50WX8, BioRad) in C18-ZipTips (Millipore)^54^. Oligosaccharides with reducing end (alditols) were separated on porous graphitized carbon columns (PGC, 5 μm particles, Hypercarb, ThermoFisher Scientific, Waltham, MA, USA) prepared in-house, with the dimensions of 10 cm (length) and 250 μm (inner diameter). The gradient, after 5 min of 98% solvent A (10 mM ammonium bicarbonate), increased solvent B (80% acetonitrile in 10 mM ammonium bicarbonate) from 2% to 45% in 41 min, then to 95% solvent B in 4 min. Solvent B (95%) was held for 5 min before re-equilibration at 98% A for 35 min. The flow rate was kept at a constant 10 μL/min using an ekspert microLC 200 HPLC system (Eksigent, AB Sciex).

The oligosaccharides were analyzed on a QTRAP 6500 ESI-triple quadrupole linear ion trap hybrid mass spectrometer (AB Sciex, Framingham, MA) in negative ion, high mass mode. A turbo V ion source was used with a 25 μm electrode. An ion spray voltage of 4200 was used. CE, DP and CXP were optimized for each transition. The final method included transitions to quantify and identify all 24 identified structures on lubricin. All transition information has been published in detail previously^35^.

### Relative quantitation and statistical analyses of MRM data

MultiQuant™ 3.0 (Sciex) was used for MRM peak integration. Area under the curve was determined for all structures and combined for total glycan abundance. Each glycan was then represented as a relative percentage of the total glycan abundance. Statistical analyses, twotailed t-tests, were carried out to compare the normal and OA synovial lubricin glycan relative abundances. GraphPad Prism 5.00 for windows (GraphPad Software, Inc, San Diego, CA, USA) was used to carry out statistical analyses and to draw graphs (specific test utilized is described in each figure legend).

### Release and analysis of oligosaccharides from bPRG4

Oligosaccharides from bPRG4 (60 μg) were released in 1.0 M NaBH4/0.1M NaOH 50°C (100 μL) overnight, followed by neutralisation with concentrated HAc, desalted with 150μl cation exchange media AG50WX8 in 100mg C18 Strata SPE column (Phenomenex). The glycans with analyzed with LC/MS and MS/MS using PGC chromatography as described above, connected to an LTQ XL ion trap mass spectrometer (ThermoFisher Scientific) as described elswhere^55^.

### Purification, digestion and LC/MS analysis of glycopeptides from rhPRG4

Aliquots of purified proteins from rhPRG4 were treated with or without glycosidases (sialidase A and/or *O*-glycanase), followed by SDS-PAGE (3-8% Tris acetate NuPage gels (Invitrogen, Stockholm, Sweden)) and tryptic digestion, or digested in-solution with either trypsin or Lys-C as described previously^37^. The peptides were desalted with C18 stagetips^56^, followed by glycopeptide enrichment using HILIC tips prepared in–house using cottonwool^37, 57^ or commercially available columns (ThermoFisher Scientific). For LC/MS analyses, C18 columns were prepared in-house and used with formic acid: acetonitrile gradients. The glycopeptides were analyzed on an LTQ Orbitrap XL mass spectrometer (ThermoFisher Scientific) equipped with CID and ETD fragmentation as described^37^.

### Purification, digestion and LC/MS analysis of glycopeptides from SF lubricin from two OA patients

SF lubricin from patients with OA was purified as described elsewhere^52^. Briefly, SF (0.5 mL) was subjected to anion exchange chromatography, the acidic protein fraction was collected and precipitated in ethanol. The acidic proteins were reduced in 10 mM DTT (70°C, 1 hour) and alkylated in 50 mM iodoacetamide, 30 min at room temperature in the dark). Salts were removed and buffer exchanged to 50mM ammonium bicarbonate (Sigma-Aldrich) using 30 kDa cutoff filters (Merck Millipore, Billerica, MA), followed by lyophilization. The reduced and alkylated protein containing gel bands were digested with trypsin, and peptides extracted and desalted with StageTip C18 columns^56^. Peptides were separated using in-house packed C18 columns at 200 nL/min using a 120 min gradient of 5-40% buffer B (A: 0.1% formic acid, B:0.1% formic acid, 80% acetonitrile). The column was connected to an Easy-nLC 1000 system (Thermo Fisher Scientific), a nano-electrospray ion source and a Q-Exactive Hybrid Quadrupole-Orbitrap Mass Spectrometer (ThermoFisher Scientific). For full scan MS, the instrument was scanned *m/z* 350-2000, resolution 70000 *(m/z* 200), AGC target 1×10^6^, max IT 120 ms, dynamic exclusion was auto or 20 s. The twelve most intense peaks (charge states 2, 3, 4) were selected for fragmentation. For MS/MS, resolution was set to 17500 *(m/z* 200), AGC target to 5×10^5^, max IT 256 ms, and collision energy normalized collision energy (NCE) =30 with stepped NCE of 25%.

Peptide searches were performed using Mascot version 2.3.02 against the database Swissprot version 2017-06. Glycopeptide searches were carried out using both manual and software assisted interpretation using Byonics (version 2.6.46, Protein Metrics, San Carlos, CA) against human lubricin (Uniprot entry Q92954). For Orbitrap XL data (recombinant Lubricin), parameters were set as follows: precursor tolerance 10 ppm; fragment tolerance 0.5 ppm; enzyme: Lys-C or trypsin, cleavage before proline included; maximum two missed cleavages; fixed carbamidomethyl modification of cysteines, variable modification: oxidized methionine. For the glycopeptide searches in Byonics, settings were maximum two glycan modifications per peptide; glycan compositions: HexNAc(1), HexNAc(1)Hex(1), HexNAc(1)Hex(1)Sulf(1), HexNAc(1)NeuAc(1), HexNAc(1)Hex(1)NeuAc(1), HexNAc(1)Hex(1)NeuAc(1)Sulf(1), and HexNAc(1)Hex(1)NeuAc(2). The glycans were all defined as ‘common’. For QExactive data (native lubricin from OA patients) parameters were the same except for fragment ion tolerance was set to 10 ppm. In Byonics, the dataset was searched against seven additional glycans of core 2 type: HexNAc(2)Hex(1), HexNAc(2)Hex(1)Sulf(1), HexNAc(2)Hex(2)Sulf(1), HexNAc(2)Hex(1)NeuAc(1), HexNAc(2)Hex(2)NeuAc(1), HexNAc(2)Hex(2)NeuAc(2), HexNAc(2)Hex(2). Sulfated glycans were defined as ‘rare’, the remaining as ‘common’. Glycopeptide spectra with a score higher than 200 were considered, and validated manually. For positive identification, at least five b/y ions originating from peptide backbone sequence were required, and all major peaks in the spectra were assigned.

### Generation and purification of PSGL-1 constructs with and without core 2 glycosylation

The P-selectin glycoprotein ligand-1/mouse IgG2b (PSGL-1/mIgG2b) and core 2 glycosyltransferase (C2GnT-1) expression plasmids carrying puromycin and geneticin (G418 sulfate) drug resistance genes, respectively, were constructed as described before^58^. CHO-K1 cells (Cat no CCL-61, ATCC, Manassas) were cultured in 75-cm^2^ T-flasks (Nunc, Roskilde, Denmark) in Dulbecco’s Modified Eagle’s medium (DMEM, Lonza Group Ltd., Basel, Switzerland) supplemented with 10% fetal bovine serum (FBS; Invitrogen), 2 mM L-glutamine (Invitrogen) and 100 units/mL penicillin, and 100 μg/mL streptomycin (Invitrogen). Cells were maintained in a humidified incubator at 37°C and 5.0% CO_2_ and transiently transfected 24 hours post-seeding at a cell confluency of 70-80%. The total concentration of plasmids encoding glycosyltransferases used for each transfection of 75-cm^2^ T-flask using the Lipofectamine 2000 Transfection Reagent Kit (Invitrogen AB) was 24 μg ensuring that equal concentrations were used (12 μg/plasmid for double transfection and 8 μg/plasmid for triple transfections). In order to generate core2 structures on PSGL-1/mIgG_2b_, CHO cells in 75 cm^2^ T-flask were transfected with plasmids encoding PSGL-1/mIgG_2b_ and C2GnT-1 glycosyltransferase supporting the generation of core 2. CP-55 was generated by transfecting the cells with plasmid encoding PSGL-1/mIgG_2b_ only.

For sodium dodecyl sulfate-polyacrylamide gel electrophoresis (SDS-PAGE)/western blotting, PSGL-1/mIgG_2b_ was purified from clarified supernatants on goat anti-mIgG agarose beads (Sigma-Aldrich; 60 μL slurry per 10 ml supernatant) by rolling head over tail at 4 °C overnight. The beads with captured fusion proteins were washed three times in PBS, resuspended in 4× SDS sample buffer without reducing agent (Invitrogen), and incubated at 95°C for 5 min for protein denaturation.

### Galectin-3 binding to SF lubricin

To measure galectin-3 binding to lubricin, 1 μg/mL recombinant galectin-3 was coated on an ELISA plate and bound SF lubricin was detected by mouse monoclonal anti-lubricin antibody clone 9G3 (Merck Millipore) followed by an HRP conjugated goat anti-mouse IgG (H+L) cross-adsorbed secondary antibody (Invitrogen). The plate was washed 3-5 times between each incubation cycle, and after incubation the plate was stained with 1-Step^™^ Ultra TMB-ELISA Substrate Solution and absorbance was read at 450 nm.

### Lubricin quantification in SF

Lubricin was quantified using an in-house sandwich ELISA method. 96-well Nunc-Immuno maxisorp plates (ThermoFisher Scientific) were coated with 1 μg/mL monoclonal anti-lubricin antibody clone 1E12 (in-house antibody against rhPRG4 mucin domain, produced by Capra Science, Sweden) in PBS at 4°C over-night. After blocking with 3% BSA in TBS-Tween (TBS pH 7.4 + 0.05% Tween-20), SF samples that were pre-diluted in assay buffer (1%BSA in TBS-Tween), were loaded and incubated for 90 minutes. The captured lubricin was incubated with 2 μg/mL biotinylated Helix Aspersa Agglutinin lectin (HAA) (Sigma-Aldrich St. Louis, MO, USA) for 1 hour, followed by a 1-hour incubation with 0.2 μg/mL horseradish peroxidase (HRP) conjugated streptavidin (Vector Laboratories). The plate was washed 3 times with washing buffer (TBS-Tween) between each incubation procedure, and 5 times before staining step. After staining with 1-Step^™^ Ultra TMB-ELISA Substrate Solution (ThermoFisher Scientific). Absorbance was read at 450nm.

### Endogenous synovial galectin-3

Endogenous galectin-3 levels in OA and control SF were measured by the Galectin-3 Test ELISA kit (BG Medicine Inc. Waltham, MA, US) according to manufacturer’s instructions. OA samples were measured in duplicates and control samples were measured once due to limited amounts of material. Statistical evaluation of ELISAs were performed by GraphPad Prism 8 (GraphPad Software, LLC, USA). Significant differences were calculated by Non-parametric t test using Mann-Whitney comparison.

### Galectin-3 binding to PSGL-1 constructs analyzed by SDS-PAGE/Western blot

For sodium dodecyl sulfate-polyacrylamide gel electrophoresis (SDS-PAGE)/Western blotting, PSGL-1/mIgG_2b_ was purified from clarified supernatants on goat anti-mIgG agarose beads (Sigma-Aldrich; 60 μL slurry per 10 ml supernatant) by rolling head over tail at 4 °C overnight. The beads with captured fusion proteins were washed three times in PBS, re-suspended in 4× SDS sample buffer without reducing agent (Invitrogen), and incubated at 95°C for 5 min for protein denaturation. The samples were analysed by SDS–PAGE using 3–8% Tris-acetate gradient gels and Tris-acetate SDS running buffer (Invitrogen). Precision protein standard (Hi-Mark, Invitrogen) was applied as reference for protein molecular weight determination. Separated proteins were blotted using iBlot (Invitrogen) in combination with nitrocellulose membranes (Invitrogen). The membranes were blocked with 1x carbo-free Blocking solution (Vector laboratories, Burlingame, CA), which was also used for dilution of antibodies and lectins. A mouse anti-human CD162 was used to detect the *N*-terminal part of PSGL-1 (1:1 000 dilution; BD Pharmingen, San Diego, CA, USA) together with an HRP-conjugated polyclonal goat anti-mouse IgG (Fab-specific, 1:10 000 dilution; Sigma Aldrich) used as secondary antibody. Biotinylated Gal-3 lectin (diluted 1:100) was used to detect galectin-3 together with strepavidin (diluted 1:10 000; Sigma Aldrich). Bound antibody and lectins were visualized by chemiluminescence using the ECL kit according to the manufacturer’s instructions (GE Healthcare, Uppsala, Sweden).

### Surface plasmon resonance (SPR)

Binding of rhPRG4 and bPRG4^51^ to galectin-3 was assessed using a Biacore X100 SPR instrument (GE Healthcare, Pittsburg PA). Galectin-3 (R&D Systems Inc., Minneapolis, MN, USA) was covalently attached to the CM5 chip by running 50 ug/ml galectin-3 solution in 20 mM sodium acetate solution at pH 5 giving a response unit of 3500. Injection of lubricin was performed as described^59^. Lactose was used as a regeneration buffer after lubricin injections^60^.

## Acknowledgements

This study was funded by grants for the Swedish state under the agreement between the Swedish government and the county council, the ALF-agreement (ALFGBG-722391 and ALFGBG-726801), the Swedish Research Council (621-2013-5895 and 2018-03077), Kung Gustav V:s 80-års foundation, Petrus and Augusta Hedlund’s foundation (M-2016-0353) and AFA insurance research fund (dnr 150150). Hakon Leffler at Lund University is gratefully acknowledged for the gift of recombinant galectin-3. Sofia Grindberg and Paula-Therese Kelly Pettersson at Danderyds Hospital and Lotta Falkendahl at University of Gothenburg are acknowledged for their assistance in collecting samples. Thanks to the Proteomics Core facility Gothenburg University for access to Byonics software.

## Competing interests

GJ, RK and TS authored patents related to rhPRG4 and hold equity in Lubris BioPharma LLC. TS is also a paid consultant for Lubris BioPharma, LLC. SF and NGK authored a patent using lubricin for diagnostics. KT, LA, SH, YM, SR, JH, OR, RB, MS, AK and TE declare no competing interest.

## Ethical approval

All individuals gave written consent and the procedure was approved by the Conjoint Health Research Ethics Board of the University of Calgary (REB15–0880) or by the Ethics Committee at Sahlgrenska University Hospital (ethical application 172-15). All methods were performed in accordance with the relevant guidelines and regulations.

## Notes

#### Summary of Updates

Synovial fluid lubricin (proteoglycan 4) is a mucin-type O-linked glycosylated (60% of the mass) biological lubricant involved in osteoarthritis (OA) development. Lubricin has been reported to be cross-linked by synovial galectin-3 on the lubricating articular surface. Here, we confirm that binding to galectin-3 depended on core-2 O-linked glycans, where surface plasmon resonance of a recombinant lubricin (rhPRG4) devoid of core-2 structures lacked binding capacity to recombinant galectin-3. Both galectin-3 levels and interactions with synovial lubricin were found to be decreased in late-stage OA patients coinciding with an increase of truncated and less sialylated core 1 O-glycans. These data suggest a defect cross-linking of surface active molecules in OA and provides novel insights into OA molecular pathology.

