## Supplemental figures for "Core-2 *O*-glycans are required for galectin-3 interaction with the osteoarthritis related protein lubricin"

**Supplementary Figure 1**. *O*-glycosylated amino acids on recombinant lubricin (rhPRG4) were identified with LC/MS and ETD fragmentation of the tryptic/Lys-C peptide digest. Glycosylated sites are highlighted in grey. Glycosylation sites found on human synovial lubricin from analyses of pooled and purified lubricin from RA and OA patients (Ali et al 2014) are underlined.

```
     MAWKTLPIYL LLLLSVFVIQ QVSSQDLSSC AGRCGEGYSR DATCNCDYNC QHYMECCPDF
61   KRVCTAELSC KGRCFESFER GRECDCDAQC KKYDKCCPDY ESFCAEVHNP TSPPSSKKAP
121  PPSGASQTIK STTKRSPKPP NKKKTKKVIE SEEITEEHSV SENQESSSSS SSSSSSSTIR
181  KIKSSKNSAA NRELQKKLKV KDNKKNRTKK KPTPKPPVVD EAGSGLDNGD FKVTTPDTSI
241  TQHNKVSTSP KITTAKPINP RPSLPPNSDT SKETSLTVNK ETTVETKETT TTNKQTSTDG
301  KEKTTSAKET QSIEKTSAKD LAPTSKVLAK PTPKAETTTK GPALTTPKEP TPTTPKEPAS
361  TTPKEPTPTT IKSAPTTPKE PAPTTTKSAP TTPKEPAPTT TKEPAPTTPK EPAPTTTKEP
421  APTTTKSAPT TPKEPAPTTP KKPAPTTPKE PAPTTPKEPT PTTPKEPAPT TKEPAPTTPK
481  EPAPTAPKKP APTTPKEPAP TTPKEPAPTT TKEPSPTTPK EPAPTTTKSA PTTTKEPAPT
541  TTKSAPTTPK EPSPTTTKEP APTTPKEPAP TTPKKPAPTT PKEPAPTTPK EPAPTTTKKP
601  APTTPKEPAP TTPKETAPTT PKKLTPTTPE KLAPTTPEKP APTTPEELAP TTPEEPTPTT
661  PEEPAPTTPK AAAPNTPKEP APTTPKEPAP TTPKEPAPTT PKETAPTTPK GTAPTTLKEP
721  APTTPKKPAP KELAPTTTKE PTSTTCDKPA PTTPKGTAPT TPKEPAPTTP KEPAPTTPKG
781  TAPTTLKEPA PTTPKKPAPK ELAPTTTKGP TSTTSDKPAP TTPKETAPTT PKEPAPTTPK
841  KPAPTTPETP PPTTSEVSTP TTTKEPTTIH KSPDESTPEL SAEPTPKALE NSPKEPGVPT
901  TKTPAATKPE MTTTAKDKTT ERDLRTTPET TTAAPKMTKE TATTTEKTTE SKITATTTQV
961  TSITTQDITP FKITTLKTTT LAPKVTTTKK TITTTEIMNK PEETAKPKDR ATNSKATTPK
1021 PQKPTKAPKK PTSTKKPKTM PRVRKPKTTP TPRKMTSTMP ELNPTSRIAE AMLQTTTRPN
1081 QTPNSKLVEV NPKSEDAGGA EGETPHMLLR PHVFMPEVTP DMDYLPRVPN QGIIINPMLS
1141 DETNICNGKP VDGLTTLRNG TLVAFRGHYF WMLSPFSPPS PARRITEVWG IPSPIDTVFT
1201 RCNCEGKTFF FKDSQYWRFT NDIKDAGYPK PIFKGFGGLT GQIVAALSTA KYKNWPESVY
1261 FFKRGGSIQQ YIYKQEPVQK CPGRRPALNY PVYGETTQVR RRRFERAIGP SQTHTIRIQY
1321 SPARLAYQDK GVLHNEVKVS ILWRGLPNVV TSAISLPNIR KPDGYDYYAF SKDQYYNIDV
1381 PSRTARAITT RSGQTLSKVW YNCP
```

Repeat region (348-855)

**Supplementary Figure 2**

HCD spectra of a Lubricin-derived *N*-glycopeptide from a patient with osteoarthritis, analyzed with LC/MS/MS using a Qexactive instrument. Analytical conditions are described in Materials and Methods

NGTLVAFR (Asn1159)
Obs. *m/z* 1047.460 (2+)
$[M+H]^+$ = 2093.913
Glycan: 1216.423 Da ($HexNAc_2Hex_5$)
Calc. $[M+H]^+$ = 2093.912

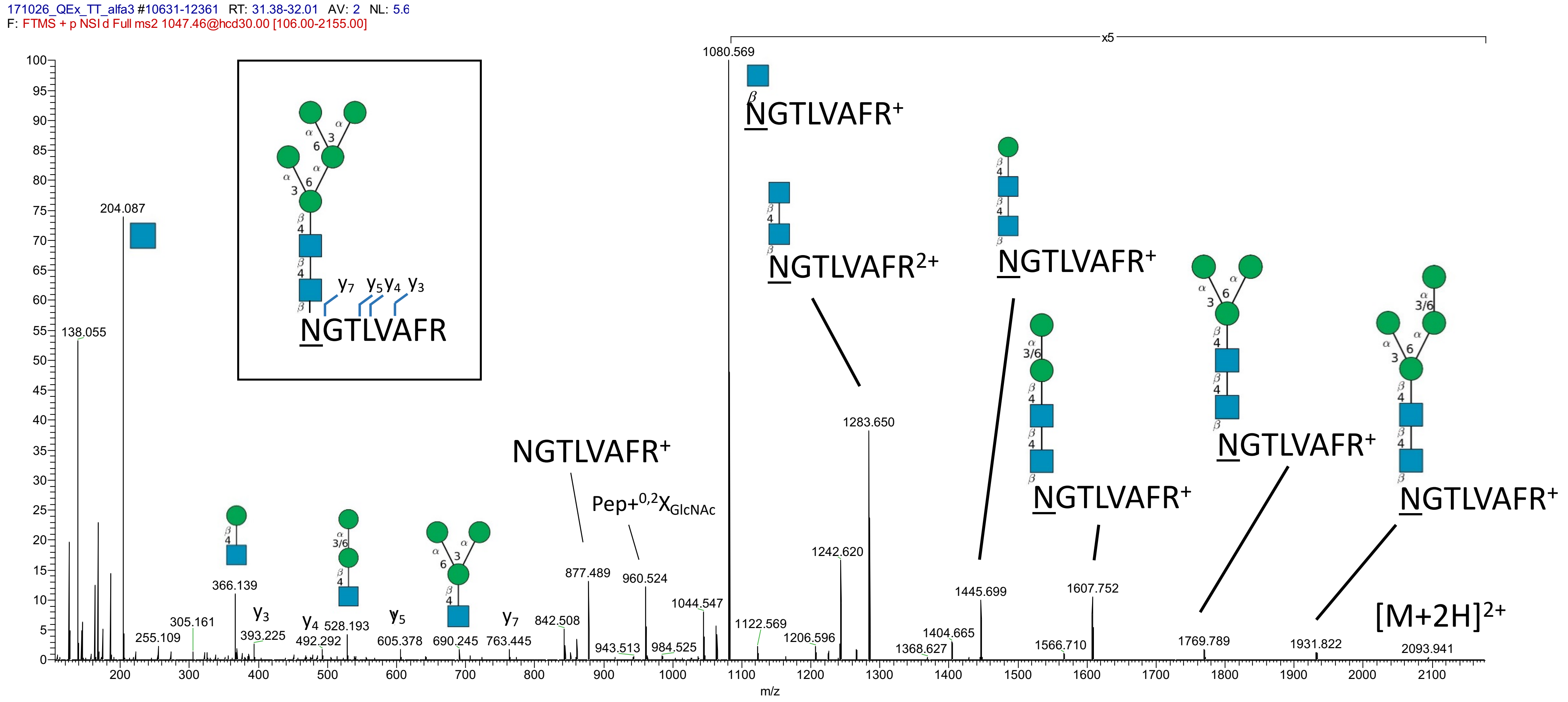

**Supplementary Figure 3**
CID spectra of a lubricin derived peptide from recombinant lubricin (rhPRG4) analyzed with LC/MS using an Orbitrap XL instrument.
Analytical conditions are described in Materials and Methods

NGTLVAFR (Asn1159)
Obs. *m/z* 1128.484 (2+)
$[M+H]^+$ = 2258.041
Glycan: 1378.476 Da ($HexNAc_2Hex_6$)
Calc. $[M+H]^+$ = 2258.045

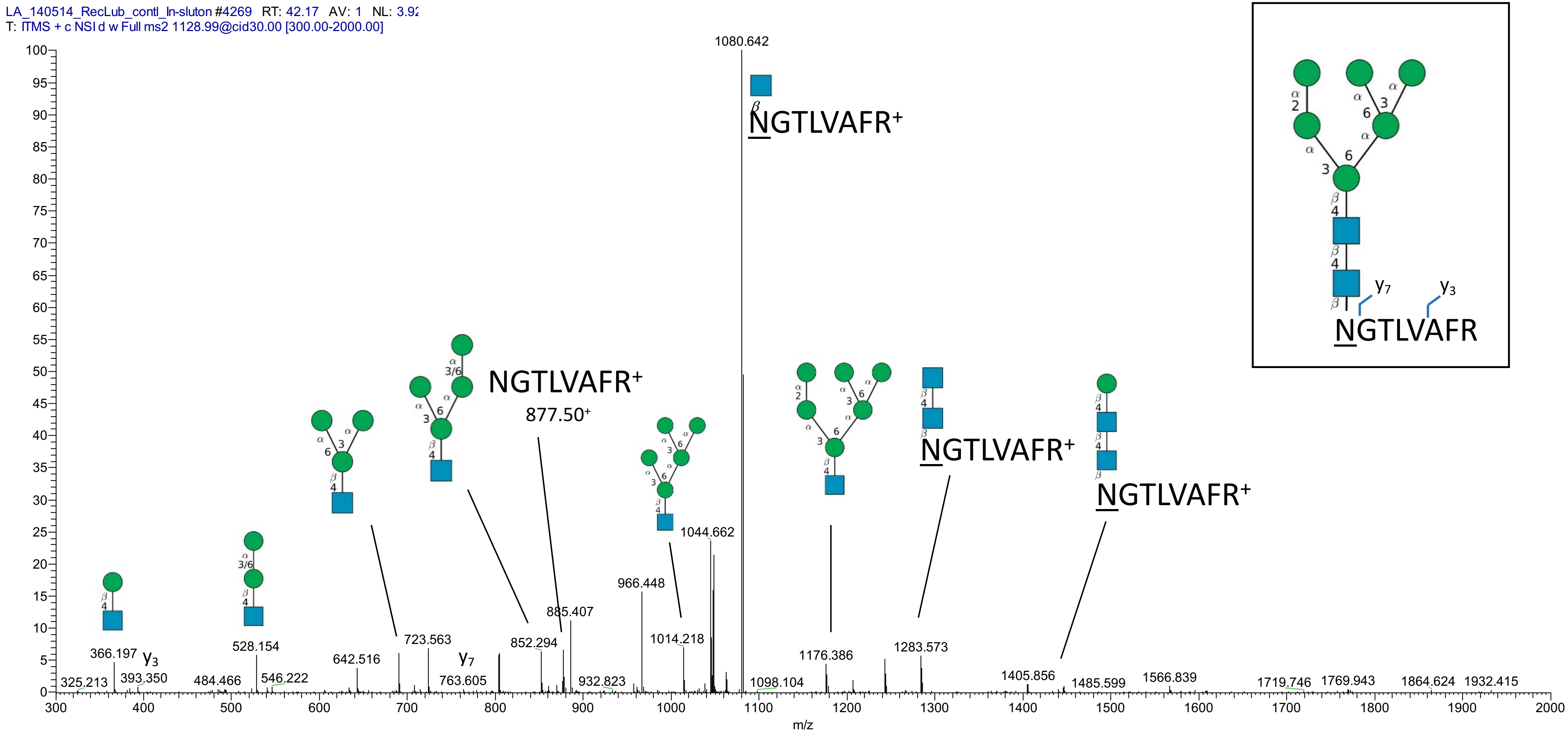
